# Combining metabarcoding and morphological approaches to identify phytoplankton taxa associated with harmful algal blooms

**DOI:** 10.1101/816926

**Authors:** Svetlana Esenkulova, Ben J.G. Sutherland, Amy Tabata, Nicola Haigh, Christopher M. Pearce, Kristina M. Miller

**Author notes:** Author for correspondence: Svetlana Esenkulova, Pacific Salmon Foundation, #300 1682 West 7^th^ Avenue, Vancouver, BC, Canada, V6J 4S6. Data Deposition: Raw sequence was uploaded to SRA under BioProject PRJNA544881 within BioSample accessions SAMN11865982-SAMN11866125.

## Abstract

Molecular techniques are expected to be highly useful in detecting taxa causing harmful algal blooms (HABs). This is the first report in Canada evaluating HABs-related species identification using a combination of morphological and molecular approaches. Microscopy, quantitative polymerase chain reaction (qPCR), and metabarcoding with multiple markers (*i.e.* 16S, 18S-dinoflagellate and 18S-diatom, large subunit (28S) rDNA) were applied on samples (n=54) containing suspected harmful algae (e.g. *Alexandrium* spp., *Chattonella* sp., *Chrysochromulina* spp., *Dictyocha* spp., *Heterosigma akashiwo*, *Protoceratium reticulatum*, *Pseudochattonella verruculosa, Pseudo-nitzschia* spp., *Pseudopedinella* sp.). Due to methodology limitations, qPCR result interpretation was limited, although good detectability occurred using previously published assays for *Alexandrium tamarense*, *H. akashiwo*, and *P. verruculosa*. Overall, the multiple-marker metabarcoding results were superior to the morphology-based methods, with the exception of taxa from the silicoflagellate group. The combined results using both 18S markers and the 28S marker together closely corresponded with morphological identification of targeted species, providing the best overall taxonomic coverage and resolution. The most numerous unique taxa were identified using the 18S-dinoflagellate amplicon, and the best resolution to the species level occurred using the 28S amplicon. Molecular techniques are therefore highly useful for HABs taxa detection, but currently depend on deploying multiple markers for metabarcoding.

## 1. Introduction

Phytoplankton form the base of the marine food web and are required to support healthy aquatic ecosystems. In some circumstances, however, high-biomass events and/or proliferation of certain algal species can cause harm to aquatic animals through a variety of means including disruption of the food web, shellfish poisoning, the development of low oxygen ‘dead zones’ after bloom degradation, and fish kills through toxins, gill damage, or hypoxia (Rensel and Whyte 2004). Collectively, these events are termed harmful algal blooms (HABs). Importantly, there is a general scientific consensus that public health, fisheries, and ecosystem impacts from HABs have all increased over the past few decades (e.g. Andersen 2012; Hallegraeff 2004).

The coastal waters of British Columbia (BC), Canada, in the northeast Pacific Ocean, have one of the longest documented histories of severe HABs going back to the first reported case in 1793 (Vancouver 1798). A government program for monitoring the presence of toxins in shellfish was established in the early 1940s (Taylor and Harrison 2002) and since then paralytic shellfish poisoning (PSP) closures have occurred every year. The BC salmon aquaculture industry initiated and has been supporting the Harmful Algae Monitoring Program (HAMP) since the 1990s due to the devastating effects of harmful algae on farmed fish (Horner et al. 1997; Rensel and Whyte 2004). During 2009–2012, direct losses to the BC salmon aquaculture industry from HABs were ~13 M USD (Haigh and Esenkulova 2014). HABs are currently one of the most significant risks for the BC aquaculture industry, regularly causing severe economic losses through finfish/shellfish mortalities and shellfish harvest closures due to toxin accumulation (Whyte et al. 1997). Therefore, there is an ongoing and pressing need for monitoring and research on HABs phenomena in coastal BC.

Monitoring HABs typically depends on the effective identification and enumeration of species of concern in water samples. Algal cell identification has long been accomplished based on morphology revealed through visual microscopic examination. Although traditional light microscopy currently remains the standard, it has limitations when it comes to certain species and strains that cannot be easily visualized or cannot be discriminated between harmful and benign variants based on morphological characteristics alone (Hallegraeff 2004). Moreover, microscopic identification is highly dependent on the level of expertise and experience of the individual analyzing the samples and with fewer morphological taxonomists being trained, it is increasingly difficult to keep up with the demand. In recent years, studying HABs with molecular techniques, either in tandem with morphological methods or independently, has become increasingly popular. Quantitative real-time polymerase chain reaction (qPCR) is a powerful method for detecting and quantifying DNA over a broad dynamic range (Livak and Schmittgen 2001). This method has been used for species-specific harmful algal detection and enumeration (e.g. Antonella and Luca 2013; Eckford-Soper and Daugbjerg 2015; Scholin et al. 2011). High-throughput platforms, such as the Fluidigm BioMark, allow for the use of multiple species-specific probes for simultaneous detection and enumeration of multiple taxa (Medlin and Orozco 2017) with the potential for both time and cost savings, as well as the additional ability to identify cryptic species, compared to traditional light microscopy. Next-generation sequencing (NGS) methods, as applied through metabarcoding, allow for millions of sequencing reactions to be performed in parallel, resulting in the ability to generate massive amounts of sequencing data (Goodwin et al. 2016; Valentini et al. 2016). Sequence-based taxonomic approaches can allow for the identification of multiple species of interest, including nano- and picoplankton, rare and fragile taxa, and cryptic species, in a reproducible and cost effective manner (e.g. Eiler et al. 2013).

The objective of the present study was to obtain microalgal taxa that are known or suspected to be harmful to cultured fish and shellfish in BC and to identify these taxa through light microscopy as well as genetic methods. Fish-killing algae targeted included *Chaetoceros concavicorne*, *C. convolutus*, *Chattonella* sp., *Chrysochromulina* spp., *Cochlodinium fulvescens*, *Dictyocha* spp., *Heterosigma akashiwo*, *Karenia mikimotoi*, and *Pseudochattonella verruculosa.* Shellfish-poisoning algae included *Alexandrium* spp., *Dinophysis* spp., *Protoceratium reticulatum*, and *Pseudo-nitzschia* spp. During this study, cultures and water samples for identification of these species were primarily acquired from HABs occurring in coastal BC. Here we describe, compare, and cross validate coastal BC harmful algae identification based on morphology, qPCR, and metabarcoding.

## 2. Methods

### 2.1. Ethics Statement

No permits were required for collection of water samples in Canadian coastal waters.

### 2.2. Sample Collection

All samples used in this study were collected in coastal BC waters (Table 1), except for two that were obtained from the Provasoli-Guillard National Center for Marine Algae and Microbiota (Samples s01, s02) that were used to test primer specificity and to optimize qPCR reactions. All samples (n=54), with their respective sampling locations, are summarized in Table 1 and Fig. 1.

**Table 1.**
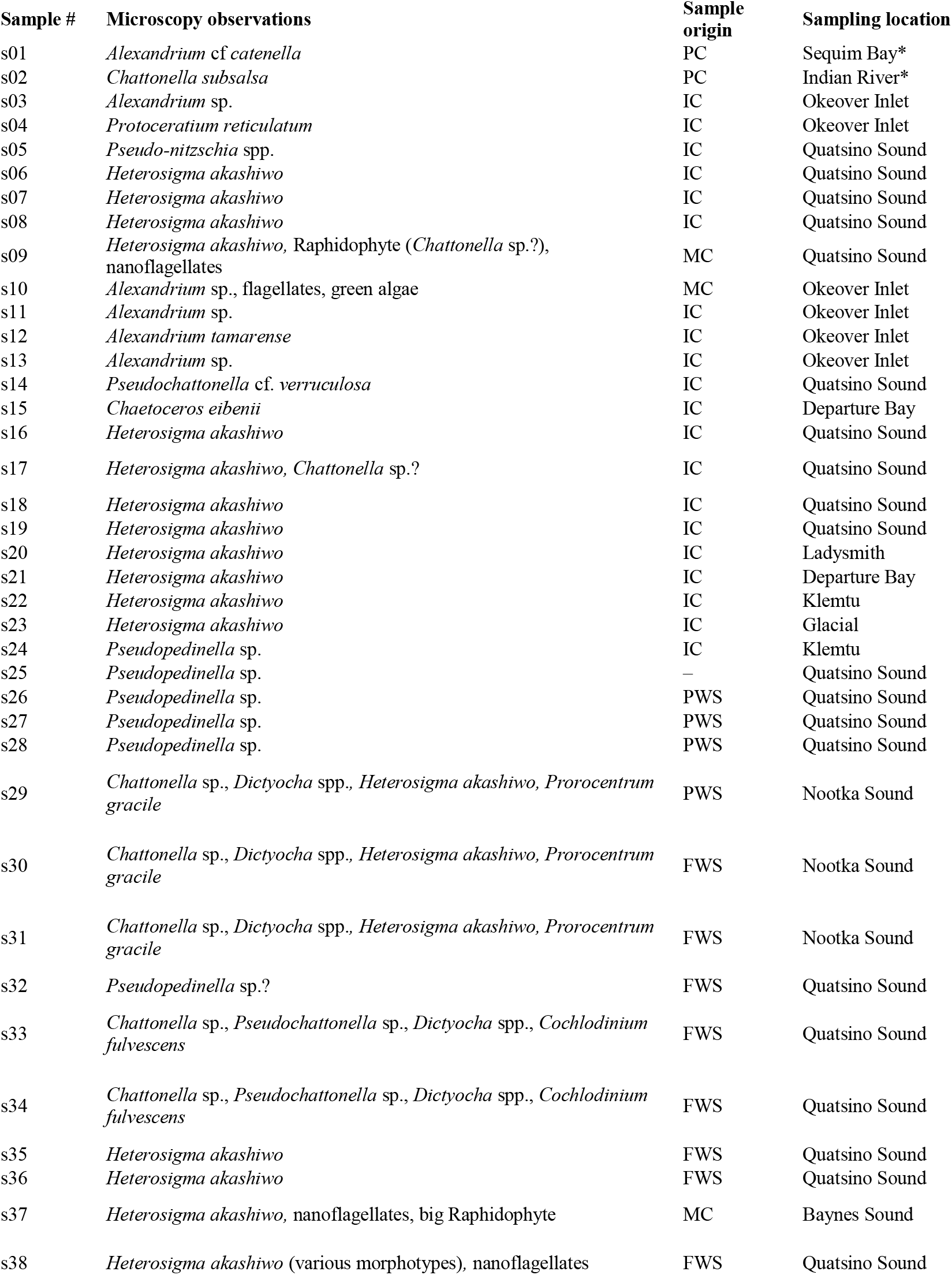
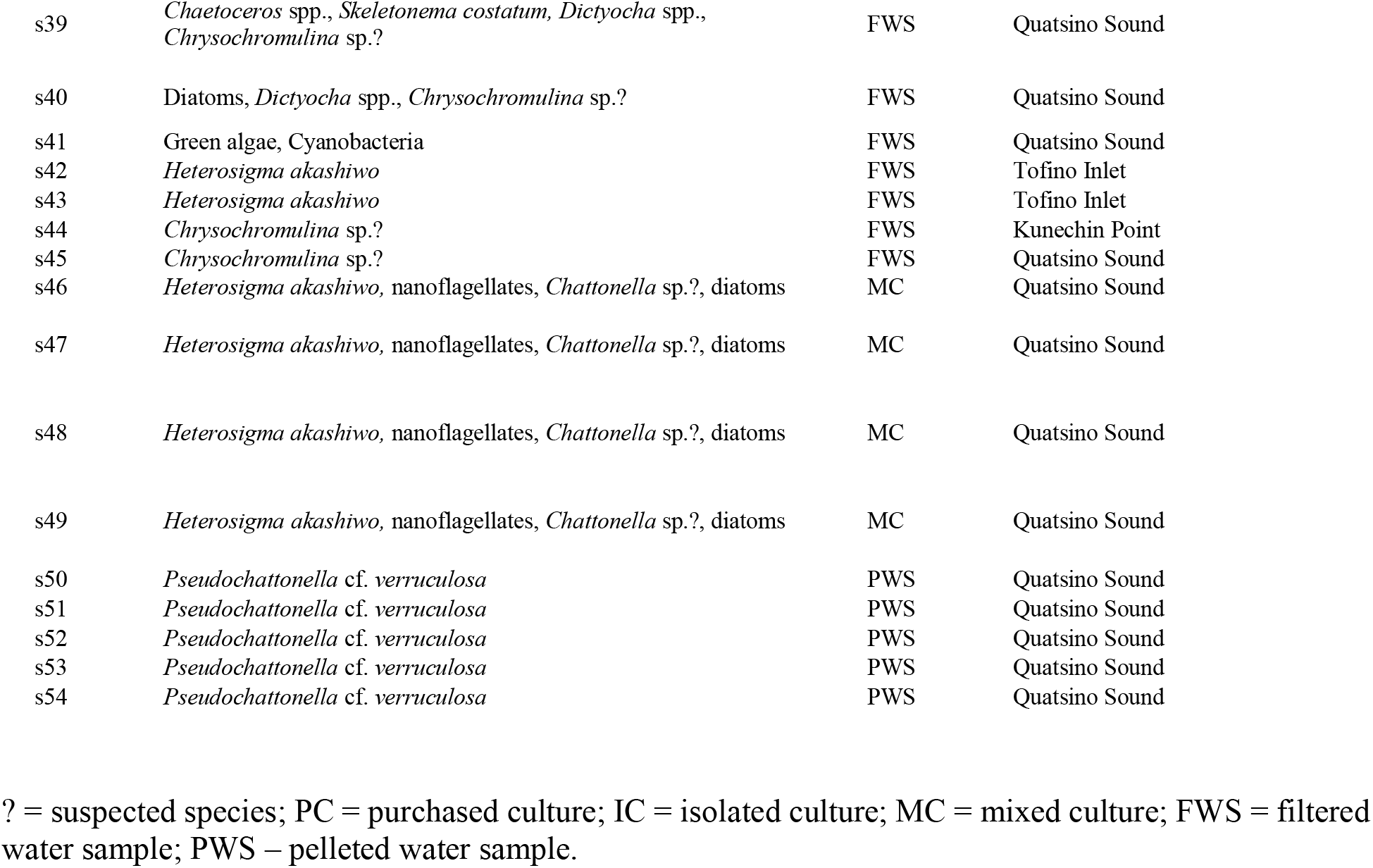
Sample identifiers, microscopy observations, and sample origin and collection location. Cultures from USA are denoted by an asterisk, with s01 being from North Pacific Ocean, Sequim Bay, Washington and s02 from North Atlantic Ocean, Indian River Bay, Delaware.

**Figure 1.**
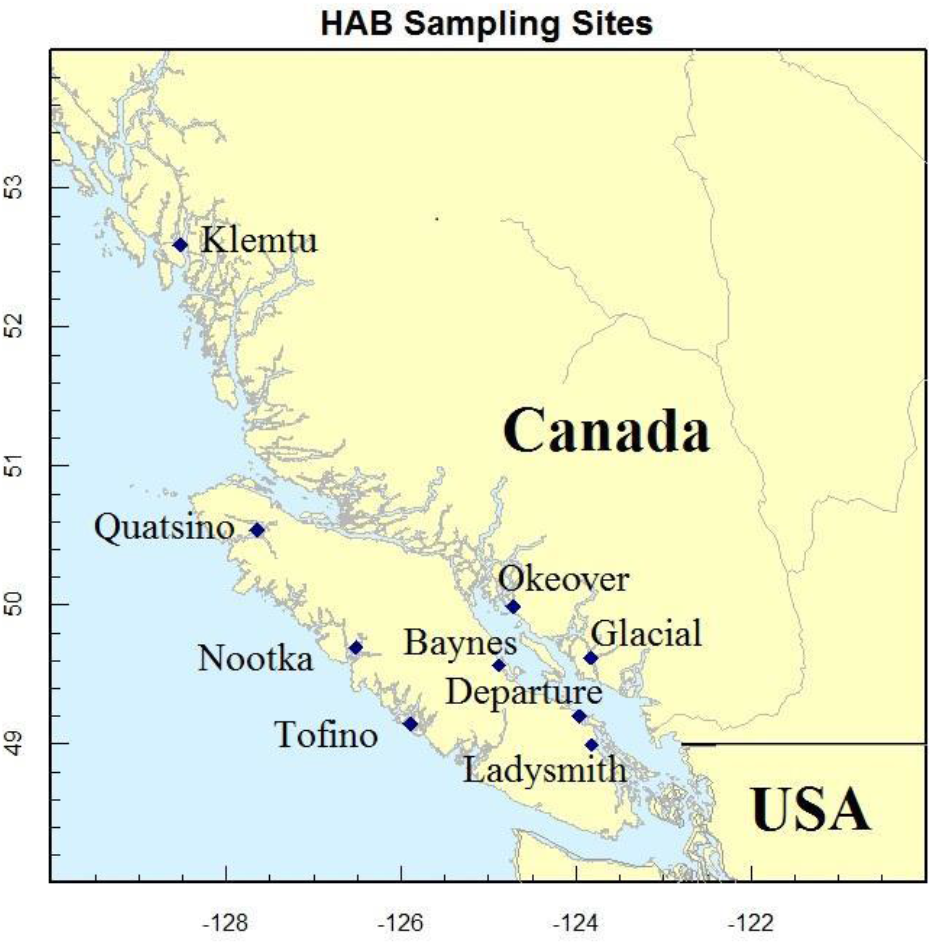
Map of Vancouver Island and British Columbia coast with locations of the sampling stations.

Field samples of algae were obtained by collecting seawater with a 1 L sampling bottle (Venrick 1978). Samples to be used for culturing and genetic analyses were kept cool and dark until processed. Samples for microscopic identification were immediately preserved with Lugol’s iodine (Andersen and Throndsen 2004) and then shipped to the laboratory for taxonomic analysis. Identification and enumeration of phytoplankton based on morphology was done with a compound microscope using a Sedgewick-Rafter slide (Guillard 1978). Identification based on morphology was done to the lowest possible taxonomic level (Hasle 1978), by an experienced phytoplankton taxonomist.

### 2.3. Cultures

Non axenic cultures for this study were isolated by serial dilution from the unpreserved field water samples and from sediments collected from sites with HABs observed in the past (see Table 1 for sample origins), as some harmful algae produce cysts that play an important role in initiating subsequent blooms (Anderson et al. 2003). Several sediment samples were taken from three areas around Vancouver Island (*i.e.* Baynes Sound, Okeover Inlet, Quatsino Sound; Fig. 1) in late winter and spring 2013. Target species for isolation and germination were *Alexandrium* spp. due to the numerous PSP closures recorded in Baynes Sound and Okeover Inlet in summer 2012 (DFO, 2012), as well as *H. akashiwo* and suspected *Chattonella* spp. that caused fish mortalities in late summer 2012 in Quatsino Sound (Haigh, personal observation). Sampling was done using van Veen and Ekman grab samplers. Sediment samples (3 – 5 L) were taken from depths of 5 – 45 m. Sites were chosen based on the assumption that cysts accumulate in the same areas as silt and clay (Wall and Dale 1967). The upper 3 cm of sediments in the grab samples were collected, mixed well, and immediately placed in containers and kept cool and dark.

Culture establishment from the collected samples occurred as described in detail previously (Esenkulova et al. 2015). In brief, sediments were sieved to produce a 20 – 120-μm fraction and incubated in enriched natural seawater medium (Harrison et al. 1980) at 18 °C under continuous illumination (20 μmol photons m^−2^ s^−1^ provided by full-spectrum fluorescent lamps). Serial dilutions were performed in a laminar flow hood using a micropipette and sterile 24-well plates. Several cultures of *A. tamarense*, *H. akashiwo*, and *Chattonella* spp.? (question mark indicates that taxon is suspected, but not positively identified by light microscopy) were successfully established as well as a few non unialgal (mixed) cultures. Isolation from water samples produced cultures of *Alexandrium* sp., *Chaetoceros eibenii* (a non-harmful species), *Chattonella* sp.?, *H. akashiwo*, *Protoceratium reticulatum*, *Pseudochattonella* cf. *verruculosa*, and *Pseudo-nitzschia* spp.

### 2.4. DNA Extraction and Purification

Water samples for genetic analyses were either filtered or pelleted. For filtered water samples, 60–100 mL of water was filtered through 25-mm or 47-mm GF/F Whatman filters and then frozen at −20 °C or stored in 75–95% ethanol until DNA extraction. A section of ¼ of the 47-mm filters or ½ of the 25-mm filters were used for the extraction. Water samples concentrated by centrifugation were subsampled and 1.5 – 40 mL centrifuged at 8,000 x g, the supernatant removed, and the resultant pellet stored in 95% ethanol at 4 °C until DNA extraction. The variation in sampling volumes and sample methods occurred due to the large variation of suggested methods in the available literature. Cultured samples for molecular analyses were taken from ~200-mL cultures in a laminar flow hood, whereby samples of well-mixed cultures were divided in 2 × 20- mL subsamples and centrifuged for 15 min at 5,000 x g for 10 min. The supernatant was decanted, samples were re-suspended in 1 mL of phosphate buffered saline, transferred to 2-mL microfuge tubes, and then centrifuged again at 4,000 g for 10 min. Finally, the supernatant was again removed and cell pellets frozen at −20 °C until extraction. Samples were frozen for less than four months.

DNA was extracted and purified from all samples using the DNeasy Blood and Tissue kit (Qiagen, Toronto, Canada) as per manufacturer’s instructions, with the addition of a homogenization step using a TissueLyser 2 homogenizer (Qiagen) with a 4-mm steel bead. DNA concentration was measured by spectrophotometry (NanoDrop, ND-1000).

### 2.5. qPCR Assays

A literature search was undertaken to identify all published (as of 2014) exclusively TaqMan-based qPCR assays for harmful algal taxa of interest in the northeast Pacific Ocean. TaqMan assays were found for five targeted taxa: six assays for *Alexandrium* spp., two for *Chattonella* spp., three for *H. akashiwo,* two for *K. mikimotoi*, and two for *P. verruculosa*. Additional published TaqMan assays for known harmful algae from other parts of the world were also included, bringing the total number of target algal taxa to 28 and the number of assays to 39 (Table S1).

The qPCR reactions using all assays (Table S1) were conducted on a Fluidigm BioMark™ platform (Fluidigm Corporation, San Francisco, CA, USA) using a 96×96 dynamic array to run 9,216 reactions simultaneously (96 samples with 96 assays) as described in detail in Miller et al. (2016). To reduce the effect of PCR inhibitors that can be problematic in algae, samples were tested at concentrations of 10 and 2.5 ng uL^−1^ or, if the DNA concentration was below that, at the highest available concentration. Each assay was run in duplicate, as were the negative controls, including an extraction control (i.e., treated through the same extraction protocol but without sample addition), and a no template control (i.e., treated as per other samples through library preparation but without sample addition) was run in duplicate. No positive controls were run (discussed below).

Briefly, a pre-amplification (STA) step of 14 cycles using dilute (50 nM in a 5-uL reaction) primer pairs of each assay with TaqMan Preamp MasterMix (Applied Biosystems, Foster City, CA, USA) was performed according to the BioMark protocol (Applied Biosystems). Unincorporated primers were removed with ExoSAP-IT (Affymetrix, Santa Clara, CA, USA) and samples were then diluted 1:5 in DNA Suspension Buffer (Teknova, Hollister, CA, USA). A 5-μL sample mix was prepared for each pre-amplified sample with TaqMan Universal Master Mix (Life Technologies Corporation, Carlsbad, CA, USA) and GE Sample Loading Reagent (Fluidigm Corporation) and a 5-μL aliquot of assay mix was prepared containing 10-μM primers and 3-μM probes for each separate TaqMan assay and each was loaded onto a pre-primed dynamic array. An IFC controller HX pressurized and mixed the assays and samples from their individual inlets on the dynamic array and the PCR was run on the BioMark with the following conditions: 50 °C for 2 min, 95 °C for 10 min, followed by 40 cycles of 95 °C for 15 s and 60 °C for 1 min. Output data was analyzed and the cycle threshold (Ct) per sample determined using Fluidigm Real Time PCR Analysis software (Fluidigm Corporation). Cycle threshold values for qPCR replicates were averaged in the final results. As no positive control or standard curve for primer efficiency testing was run, all qPCR results were considered qualitatively rather than quantitative, and result interpretation was limited.

### 2.6. Metabarcoding and Sequencing

Four primer pairs were selected for use in NGS metabarcoding from published studies with the goal of amplifying a broad range of algal taxa (see Table S1), including 16S (Eiler et al. 2013), 18S-dinoflagellate (Kohli et al. 2014), 18S-diatoms (Zimmermann et al. 2011), and 28S (Hamsher et al. 2011). Illumina adapters were incorporated onto the 5’ end of each primer for the attachment of Nextera XT Illumina indices (Illumina, Inc., San Diego, CA, USA) in the second round of PCR. Input sample DNA was normalized to 5 ng μL^−1^ when starting concentrations allowed. The primer sets were used as the initial primers in a two-step PCR process to construct and sequence four Nextera XT libraries according to the Illumina 16S Metagenomic Sequencing Library Preparation protocol (15044223, Rev. B). Library quantification, normalization, pooling, and sequencing on a MiSeq with a 600-cycle flow cell (MiSeq Sequencing Kit v3, 600 bp, Illumina, Inc.) were performed according to the manufacturer’s protocols, with the only modification being that the final, pooled library was run at a concentration of 16 pM with 10% PhiX control.

### 2.7. Bioinformatics

All sequence data was de-multiplexed using input sample barcodes during file export from the sequencer (Illumina, Inc.), resulting in a pair of fastq files for each individual sample. Quality of raw sequence fastq files was evaluated in FastQC (Andrews and Babraham Bioinformatics 2010) with results aggregated using MultiQC (Ewels et al. 2016). In general, the OBITools package (Boyer et al. 2016) was used for the analysis of the different amplicons, but each amplicon required specific inputs due to different features of the data. All bioinformatics steps are outlined in detail on GitHub (see *Data Availability*), and explained here per marker type (*i.e.* 28S also referred to as LSU in the pipeline, 18S, and 16S). For 28S and 18S, due to the larger size of amplicons and the lack of read merging for these paired-end datasets, only the forward reads were used. For 16S, paired-end data were used as most reads overlapped and therefore were able to be merged.

For 28S (single-end data), primers were removed using cutadapt (Martin 2011) and then reads were formatted for the OBITools pipeline using *ngsfilter* (OBITools) on each sample without any input barcodes. Each formatted input sample file was then annotated with a sample identifier in read headers with *obiannotate*. Sample labeled fastq files were then combined to a single file, with reads cut to a uniform size (*i.e*. 230 bp) to reduce singleton records due to slight differences in read length for single-end data. Subsequently, this file was moved into the standard OBITools pipeline (see below).

For 18S (single-end data), there were two different amplicons, one for dinoflagellates and one for diatoms (*i.e.* 18S-dinoflagellate, 18S-diatom). These two amplicon types were de-multiplexed using *ngsfilter*, which output a single fastq file per sample, but from both amplicon types. A custom script separated the two types of amplicons (see *Data Availability*). Obiannotate was used to annotate each sample-amplicon type with sample and amplicon identifiers. All data was then merged into a single file, cut to a uniform size using cutadapt (as above), and then moved into the standard OBITools pipeline.

For 16S (paired-end data), primers were removed using cutadapt, overlapping reads were merged using *illuminapairedend* retaining merged data with an overlap score ≥ 40 (OBITools), samples were annotated with a sample identifier, and then combined to a single file and moved into the standard OBITools pipeline.

Once all data were input in a uniform format into the main pipeline as described above, the OBITools package was used to retain a single representative accession per unique amplicon sequence, keeping record of the number of reads per sample for the accession in the accession header using *obiuniq.* Subsequently, the data were de-noised using a size filter and a low count threshold (*obigrep*) and by removing ‘internal’ sequences (*i.e.* probable PCR/sequencing errors; *obiclean*), as per standard OBITools approaches (Boyer et al. 2016). The counts per representative unique amplicon were exported using *obitab*.

The unique amplicon file for each amplicon type was annotated using BLAST (Altschul et al. 1997) run in parallel (Tange 2011), receiving ten alignments per BLAST query. For each unique amplicon accession, taxa were assigned using MEGAN (Huson et al. 2016) using the Lowest Common Ancestor (LCA) algorithm with the following parameters: min score = 100; max expected = 10^−9^; min % ID = 97; top % 10; min support % (off); and min support = 1. Any amplicon that received a BLAST result, but was not assigned using MEGAN due to parameters set within MEGAN was put in the ‘Not assigned’ category. Any amplicon without a BLAST result at all was put in the ‘Unknown’ category. Read counts were connected to annotations from the MEGAN output using custom R (R Core Team, 2018) scripts (see *Data Availability*). A threshold of at least 10 reads per sample-taxon combination was applied to reduce the potential for false positive detection and any sample-taxon combination with fewer than this was transformed to 0. Taxa proportions were calculated by dividing the count per taxon for a sample by the total number of reads for the sample. Taxon ranks and the classification of identified taxa was enabled using *taxize* (Chamberlain and Szöcs 2013; Scott et al. 2019) in R, and a custom database was created to contain assembly taxonomy, read count, and sample information. Bar plots were constructed using ggplot2 in R (Wickham 2016) and pie charts constructed using Krona (Ondov et al. 2011).

## 3. Results

### 3.1. Morphological Identification

In total, 54 samples containing either cultured (n=30) or field (n=24) collections of ten suspected harmful algal taxa identified by morphology were obtained during this study, including *Alexandrium* spp., *Chattonella* sp., *Chrysochromulina* spp., *Cochlodinium fulvescens*, *Dictyocha* spp., *Heterosigma akashiwo*, *Protoceratium reticulatum*, *Pseudochattonella verruculosa*, *Pseudo-nitzschia* spp., and *Pseudopedinella* sp. (note: *Pseudopedinella* was not initially targeted, but was acquired opportunistically at a suspected HAB event). Although other species were initially targeted, they were unable to be acquired for complete analysis, due to either low cell concentrations in field samples (*i.e. C. concavicornis*, *C*. *convolutus,* and *Dinophysis* spp.) or due to a general absence in field samples (*i.e. K. mikimotoi*). All samples obtained for this study are listed in Table 1.

### 3.2. Extraction Efficiency for Molecular Methods

Sample collection volumes, types, and preservation methods varied throughout the collection time period as sampling and extraction protocols were still being optimized. As such, DNA quantity in extractions was highly variable due to the variety in the sample state, density, and preservation methods applied. Cultured samples gave the highest yields. Filtered field samples generally resulted in lower, but adequate yields. Pelleted field samples resulted in low and often unusable yields. After extractions, 46 of the 54 samples had sufficient high-quality DNA for molecular analyses. Samples with insufficient DNA (*i.e.* s025, s026, s027, s029, s039, s051, s052, s054) were not run on qPCR, but two of these samples (*i.e.* s029 and s054) were run through metabarcoding NGS and returned OTUs (<25 and <1,350, respectively). Five samples (*i.e.* s034, s045, s046, s050, s053) had low DNA concentration, but returned positive results with qPCR and NGS.

### 3.3. Taxa Identified by qPCR

A total of 46 samples with sufficient DNA concentrations were run using qPCR. However, only seven of 39 applied TaqMan assays provided amplification with one or more of the samples (Table S2), and the rest (n=32 assays) did not return positive/suspected results in any samples. These seven assays amplified within 43 of the samples. Targeted species that were detected via qPCR included *Alexandrium* spp./*A. tamarense*, *H. akashiwo*, and *P. verruculosa*. The targeted *Chattonella* spp. had four different qPCR assays, but none resulted in detections. Other species that are potentially harmful were identified by qPCR, but were not specifically identified by microscopy. For example, cyanobacteria species were detected in nine samples and *Karlodinium micrum/veneficum* in ten samples. Nine targeted taxa were not quantifiable by qPCR due to a lack of published TaqMan assays: *C. concavicornis*, *C*. *convolutus, Chrysochromulina* spp., *Cochlodinium fulvescens*, *Dictyocha* spp., *Dinophysis* spp., *P. reticulatum*, and *Pseudo-nitzschia* spp. Due to the fact that this study ran the assays under different conditions than they were designed for, absence of positive controls, and due to algae strain variability, interpretation of qPCR results was limited. To confirm if the sample contained a target species, and therefore to determine the effectiveness of the tested assays and correspondence to microscopy, NGS was conducted. The relative effectiveness of the qPCR results in comparison to microscopy and metabarcoding is provided in the *Discussion*.

### 3.4. Metabarcoding Overview

In total, over 350 individual taxa were detected by metabarcoding NGS. NGS returned results for 48 of 48 sequenced samples. Detailed results of the analysis including all taxa and associated read counts are provided in Supplemental Tables S3-7.

The most numerous unique taxa occurred through using the 18S-dinoflagellate amplicon (Table 2), which returned 167 individual taxa, including 49 to the species level. For comparison, the 16S amplicon identified 69 individual taxa (21 to the species level), and the 18S-diatom amplicon identified 78 taxa (18 to the species level). The 28S data identified 136 individual taxa and was the most effective at resolving data to the species level (n=60 species), which is noteworthy given the importance of species-level identification when identifying harmful algae. Only one taxon, *H. akashiwo*, was identified by all four amplicons. Many similar taxa (n=20) were detected by 18S-diatom, 18S-dinoflagellate, and 28S amplicons, but only three were identified to the species level.

**Table 2.**
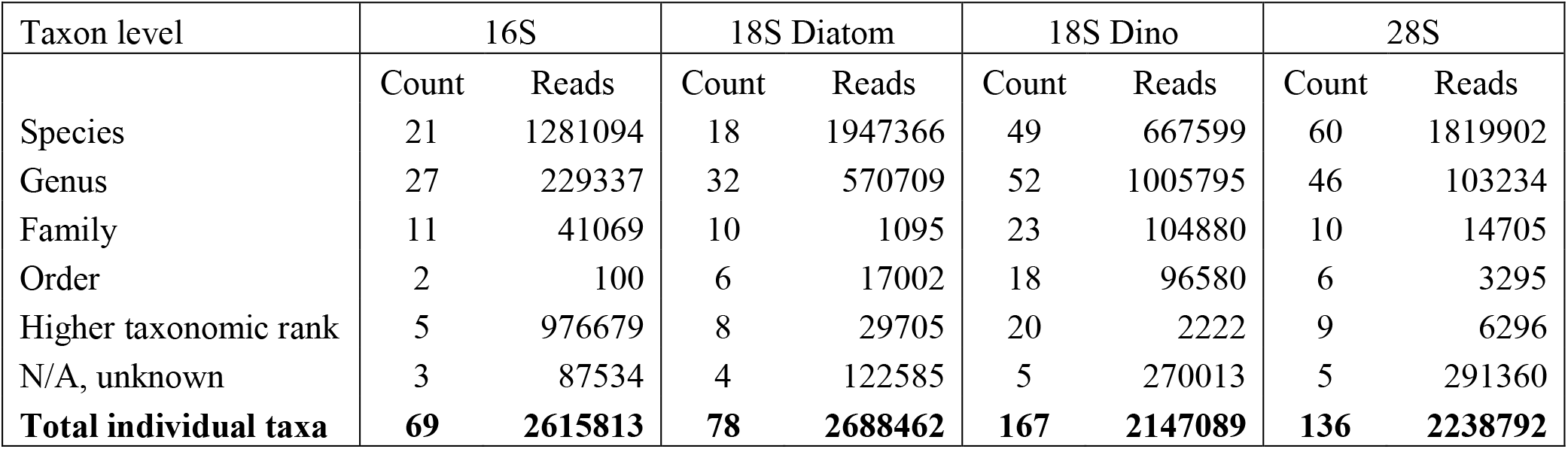
Taxon levels and number of reads detected by the four different amplicons: 16S, 18S-Diatom, 18S-Dinoflagellate, and large subunit (28S).

| Taxon level | 16S |  | 18S Diatom |  | 18S Dino |  | 28S |  |
| --- | --- | --- | --- | --- | --- | --- | --- | --- |
|  | Count | Reads | Count | Reads | Count | Reads | Count | Reads |
| Species | 21 | 1281094 | 18 | 1947366 | 49 | 667599 | 60 | 1819902 |
| Genus | 27 | 229337 | 32 | 570709 | 52 | 1005795 | 46 | 103234 |
| Family | 11 | 41069 | 10 | 1095 | 23 | 104880 | 10 | 14705 |
| Order | 2 | 100 | 6 | 17002 | 18 | 96580 | 6 | 3295 |
| Higher taxonomic rank | 5 | 976679 | 8 | 29705 | 20 | 2222 | 9 | 6296 |
| N/A, unknown | 3 | 87534 | 4 | 122585 | 5 | 270013 | 5 | 291360 |
| **Total individual taxa** | **69** | **2615813** | **78** | **2688462** | **167** | **2147089** | **136** | **2238792** |

The numbers of reads per taxonomic category, with an emphasis on microalgal groups, are listed in Table 3. Total microalgae reads detected by 18S-diatom, 18S-dinoflagellate, and 28S runs were more than 85% of all reads (not including unknowns) for each amplicon, whereas microalgae reads from 16S run comprised less than 50%. The majority of microalgal reads for all amplicons belonged to raphidophytes. The 18S-dinoflagellate amplicon detected the most microalgal groups and the 16S amplicon detected the fewest microalgal groups. Silicoflagellates (Dictyochophyceae), an important group that includes several harmful and potentially harmful species, were detected only by 18S-diatom and 18S-dinoflagellate amplicons.

**Table 3.**
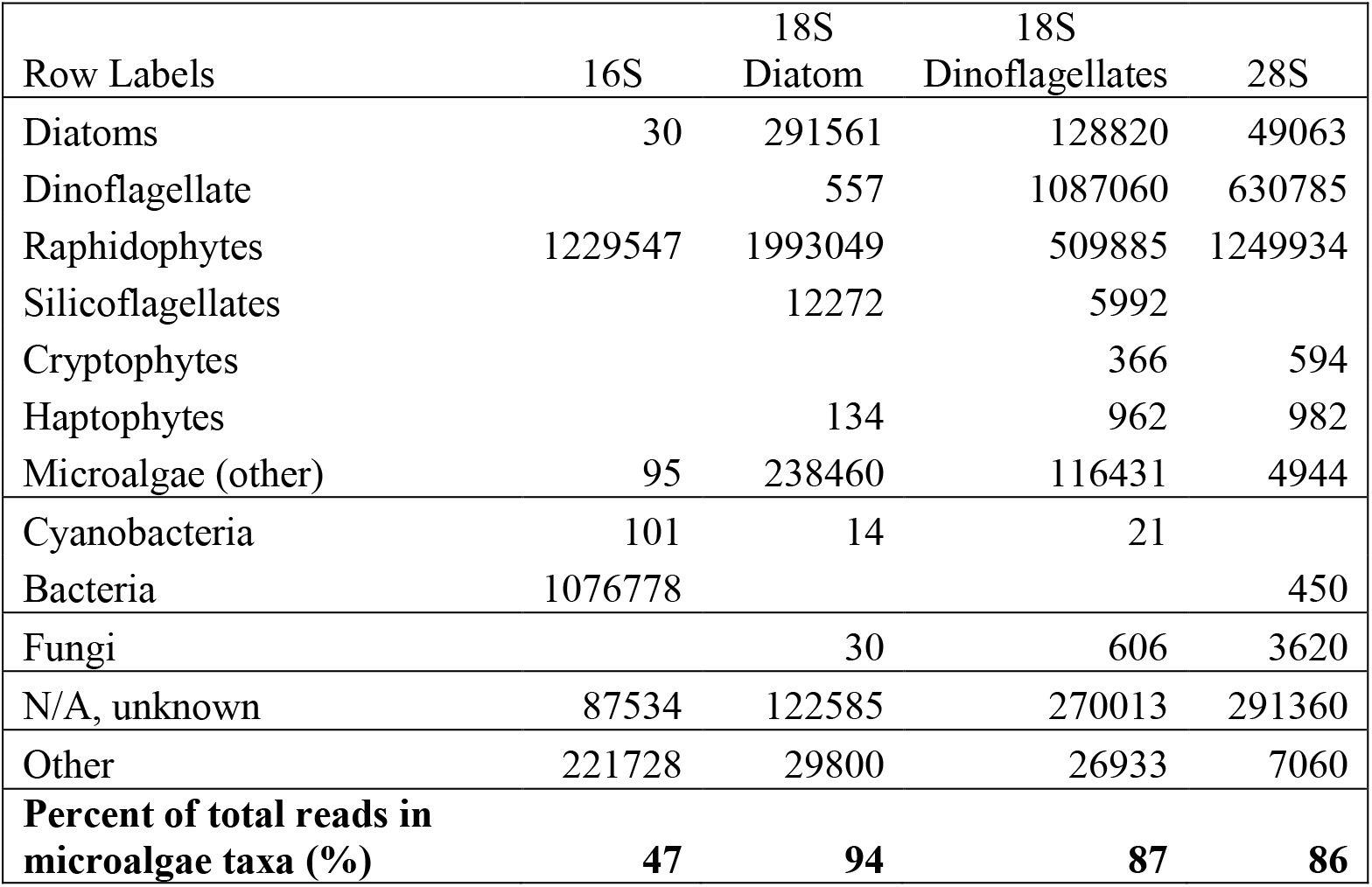
Number of reads per taxonomic category and percent of total microalgae reads.

All harmful and potentially harmful algal taxa reads (*i.e.* at the species, genus, and family level) detected by all amplicons are provided in Table 4. For a more convenient comparison between results employing different amplicons, grouping of harmful and potentially harmful algae species to genus level was done for *Alexandrium* spp., *Chattonella* spp., *Dinophysis* spp., *Karlodinium* spp., *Phalacroma* spp., *Prymnesium* spp., *Pseudo-nitzschia* spp., and *Pseudochattonella* spp.

**Table 4.** Number of reads for all detected harmful and potentially harmful algae species, genera, and families.

| Taxonomic group | 16S | 18S<br>Diatom | 18S<br>Dino | 28S | Amplicon<br>returning<br>maximum read |
| --- | --- | --- | --- | --- | --- |
| *Alexandrium andersonii* |  |  | 97 |  | 18S Diatom |
| *Alexandrium fundyense* |  |  |  | 549437 | 28S |
| *Alexandrium* spp. |  | 509 | 920011 | 14871 | 18S Dino |
| *Alexandrium tamarense* |  |  | 4300 | 292 | 18S Dino |
| **Total *Alexandrium* spp.** | **0** | **509** | **924408** | **564600** | **18S Dino** |
| *Chattonella* spp. |  | 152203 | 40 | 64380 | 18S Diatom |
| *Chattonella subsalsa* |  |  | 58 | 28 | 18S Dino |
| **Total *Chattonella* spp.** | **0** | **152203** | **98** | **64408** | **18S Diatom** |
| *Dinophysis parvula* |  |  |  | 13 | 28S |
| *Dinophysis* spp. |  |  | 119 | 111 | 18S Dino |
| *Phalacroma rapa* |  |  |  | 44 | 28S |
| **Total *Dinophysis* and *Phalacroma* spp.** | **0** | **0** | **119** | **168** | **28S** |
| *Karlodinium* spp. |  |  | 21 | 50 | 28S |
| *Karlodinium veneficum* |  |  |  | 27 | 28S |
| **Total *Karlodinium* spp.** | **0** | **0** | **21** | **77** | **28S** |
| *Prymnesium kappa* |  |  |  | 20 | 28S |
| *Prymnesium* spp. |  |  | 10 |  | 18S Dino |
| **Total *Prymnesium* spp.** | **0** | **0** | **10** | **20** | **28S** |
| *Pseudo-nitzschia* sp. B HAB-2017 |  |  |  | 33 | 28S |
| *Pseudo-nitzschia* spp. |  | 72978 |  | 11254 | 18S Diatom |
| **Total *Pseudo-nitzschia* spp.** | **0** | **72978** | **0** | **11287** | **18S Diatom** |
| *Pseudochattonella farcimen* |  | 167 |  |  | 18S Diatom |
| *Pseudochattonella* spp. |  | 1944 | 2104 |  | 18S Dino |
| **Total *Pseudochattonella* spp.** | **0** | **2111** | **2104** | **0** | **18S Dino** |
| *Chattonellaceae* |  |  | 38486 |  | 18S Dino |
| *Chrysochromulina* spp. |  |  |  | 220 | 28S |
| *Cochlodinium* spp. |  |  |  | 1016 | 28S |
| *Heterosigma akashiwo* | 1229451 | 1840799 | 471269 | 1185484 | 18S Diatom |
| *Protoceratium reticulatum* |  |  | 145010 | 43109 | 18S Dino |
| *Prymnesiaceae* |  |  | 11 |  | 18S Dino |

Overall, most of the reads of harmful and potentially harmful algae were detected by 28S and 18S-dinoflagellate amplicons (Table 4). At the species and genus levels, the 28S amplicon provided the most reads for dinoflagellate species within the *Cochlodinium*, *Dinophysis*, *Karlodinium*, and *Phalacroma* genera, as well as haptophytes in the *Chrysochromulina* and *Prymnesium* genera. The 18S-dinoflagellate amplicon detected the most reads for species within the *Alexandrium*, *Pseudochattonella*, and *P. reticulatum* genera, whereas the 18S-diatom amplicon detected the most reads for *Pseudo-nitzschia* spp. as well as the raphidophytes *Chattonella* sp. and *H. akashiwo*. There were no instances when 16S detected the most reads for any of the listed taxa.

The total reads and percentages per sample detected by different amplicons are shown in Figure 2. Specifically in this plot, when a taxon of harmful or potentially harmful algae had less than 100 reads, it was grouped within an appropriate, larger algae category, e.g. *Karlodinium* spp. with 77 reads were included in the dinoflagellates counts; *Prymnesium* spp. with 20 reads and Prymnesiaceae with 11 reads were included in haptophytes (Figure 2). A comparison of microscopy and NGS taxonomic identification in each of the 48 samples revealed that the majority of the taxa detected by microscopy also were identified in the NGS results (Supplemental Table S7). Species and genera that were positively identified by both microscopy and NGS included: *Alexandrium*, *Chaetoceros, Cochlodinium, H. akashiwo, P. reticulatum, Pseudochattonella*, and *Pseudo-nitzschia*. There were a few notable exceptions, with the two most consistently observed mismatches being (i) all suspected *Chattonella* spp. (10 field and cultures) were identified by NGS as *H. akashiwo*; and (ii) *Dictyocha* spp., *P. verruculosa*, *Pseudochattonella* sp., and *Pseudopedinella* spp. (*i.e.* all three silicoflagellate genera) were not detected in 11 out of 15 samples by any of the amplicons. Although these samples had unknown and unassigned reads in NGS results (Supplemental Table S7), the sequences for all of the species not identified by NGS but identified with microscopy were in fact present in the GenBank database, and therefore the reason for the lack of these taxa, as well as the identity of these unknown reads remains unknown.

**Figure 2.**
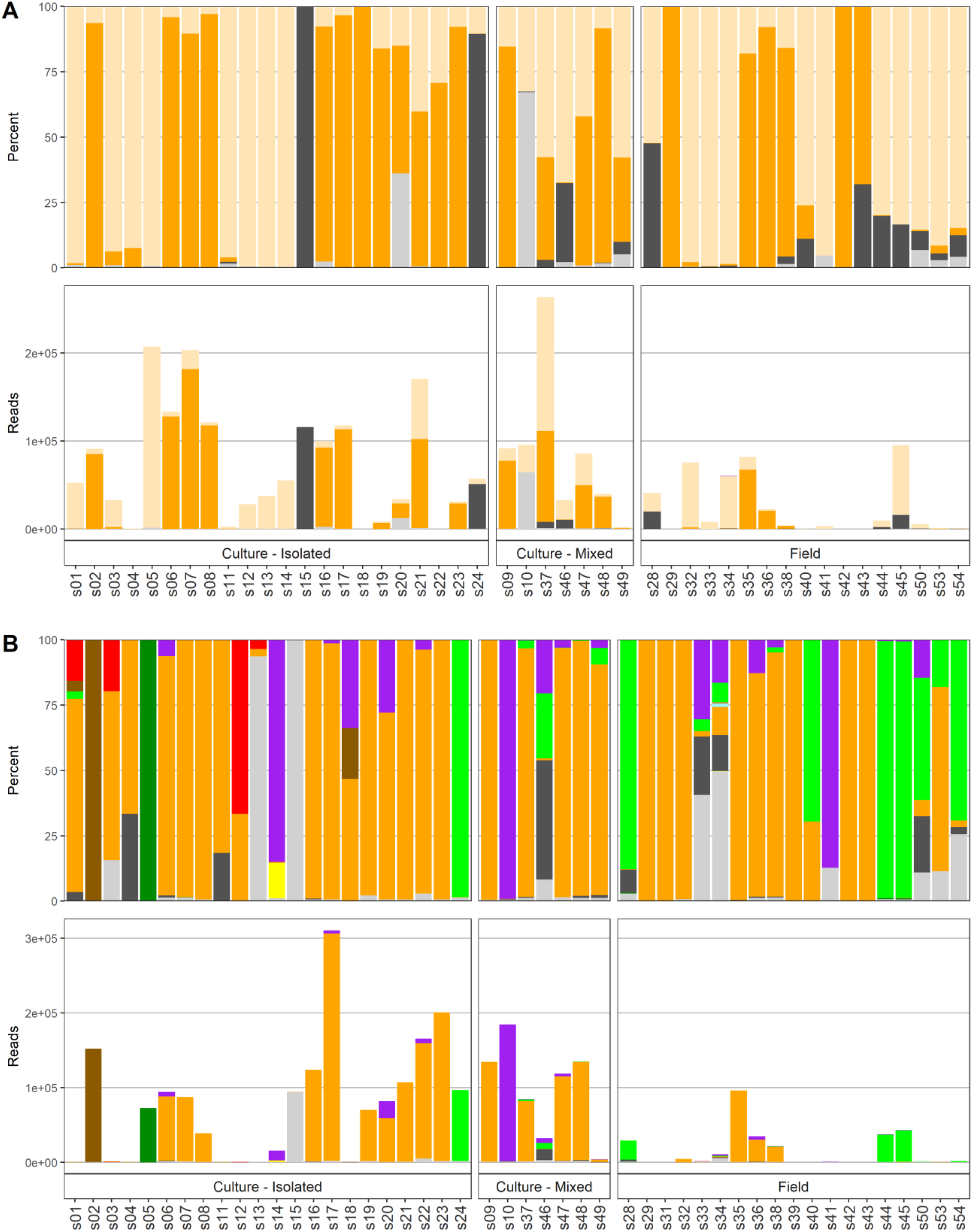
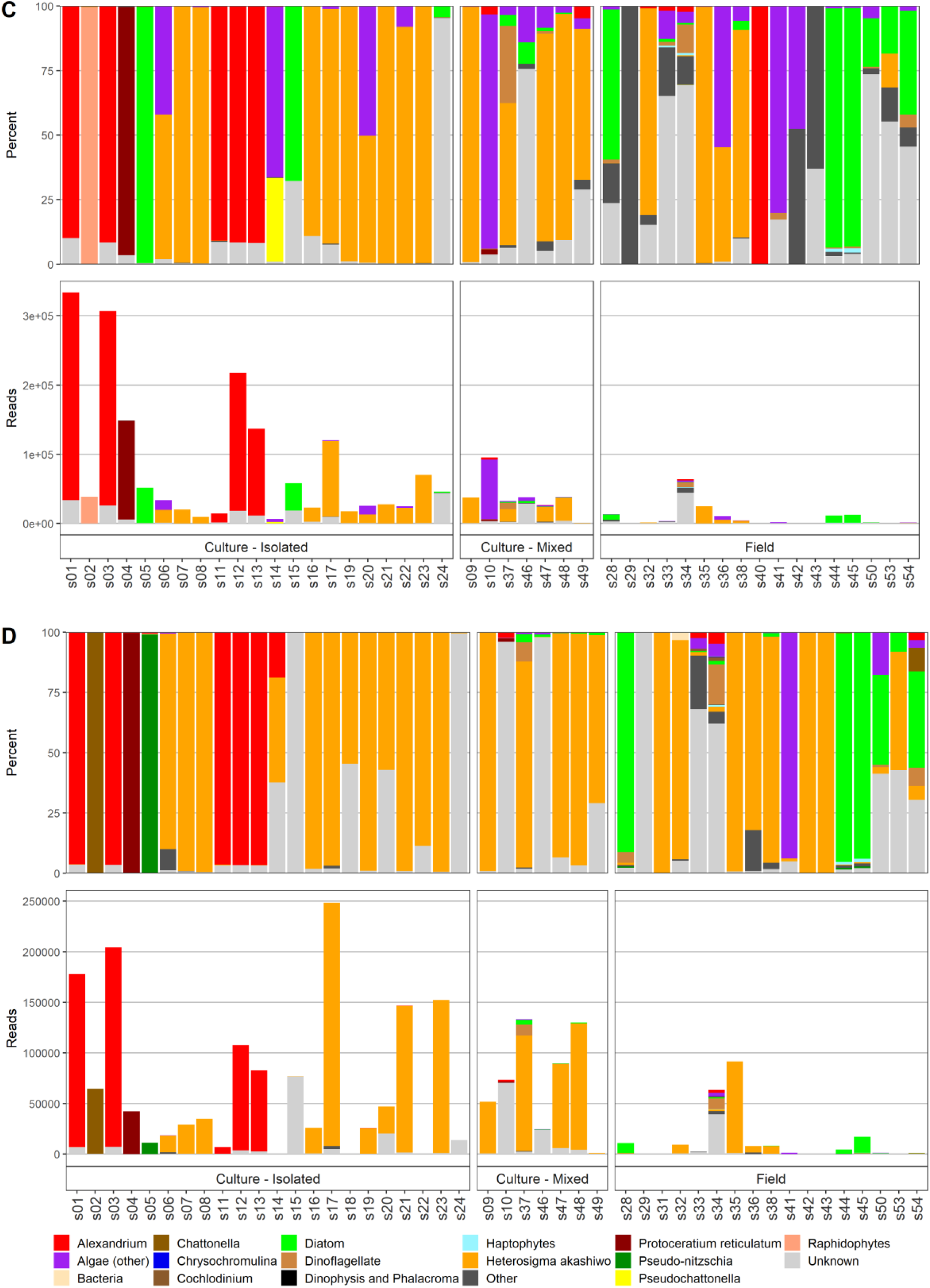
Percentages and number of reads per sample with emphasis on harmful and potentially harmful algal species: (A) 16S, (B) 18S-Diatom, (C) 18S-Dinoflagellate, and (D) 28S.

## 4. Discussion

### 4.1. Harmful Algae: Comparing Microscopy and Molecular Approaches

The term ‘algae’ comprises a diverse, polyphyletic group that encompasses organisms from widely different taxonomic domains and includes eukaryotes and prokaryotes. Major algal groups include diatoms, dinoflagellates, raphidophytes, silicoflagellates, haptophytes, and cyanobacteria. Identification methods applied to each of these groups will be outlined in detail below.

### 4.2. Diatoms

Diatoms are the major group of microalgae with approximately 250 modern genera and thousands of species (Hasle and Syvertesen 1996). There are only a few diatom taxa that are harmful due to toxin production (*i.e.* several species from the *Pseudo-nitzschia* genus) or cell structures that cause mechanical damage to gills (e.g. *C. concavicornis* and *C. convolutus*). In our study, only one sample contained a potentially harmful diatom – the culture of *Pseudo-nitzschia* sp. Positive identification of *Pseudo-nitzschia* to the species level with microscopy can be ensured only with scanning electron microscopy (SEM), which is a very costly and time-consuming technique not generally implemented in routine monitoring. In our study, both the 18S-diatom and 28S amplicons showed promising results by confirming *Pseudo-nitzschia* presence (>10,000 reads, >98% of the total reads in the cultured sample). However, it also did not discriminate to the species level. The same amplicons detected very low presence (<1%) of *Pseudo-nitzschia* spp. in five other samples where cell presence was not detected using microscopy. Most of these samples were presumed monocultures of other species that were processed on the same day (*i.e.* culture subsampled, DNA extracted and purified), and NGS results may indicate possible low-level cross contamination from high concentration, pure cultures that occurred in the lab in the same flow hood.

### 4.3. Dinoflagellates

Dinoflagellates are second to diatoms in terms of importance to marine primary production, with about 2,000 described extant species (Taylor et al. 2008). This group is particularly important for HABs, as about 75–80% of known toxic phytoplankton species belong to this group (Cembella 2003). Microscopy indicated that there were nine samples containing *Alexandrium* spp., *Cochlodinium* sp., or *P. reticulatum*. However, based on molecular techniques, additional potentially harmful dinoflagellate species were detected (*Dinophysis*, *Karlodinium*, and *Phalacroma*) and considerably higher numbers of samples containing them were identified. For dinoflagellates, as expected, the most conclusive results were obtained using the 18S-dinoflagellate and 28S amplicons.

#### 4.3.1. Alexandrium

Many species of *Alexandrium* produce toxins that cause paralytic shellfish poisoning (PSP). In BC, since the establishment of the monitoring program for toxins in shellfish in the 1940s (Taylor and Harrison 2002), PSP closures have occurred every year, and this negatively affects shellfish aquaculture and recreational harvesting. Six cultured samples of *Alexandrium* were included in this study. One was a purchased culture of *Alexandrium* cf. *catenella* and five were locally established cultures microscopically identified as *Alexandrium* sp. and *A. tamarense;* four of these cultures were monocultures and one was a mixed culture that also contained green algae and flagellates. The most effective metabarcoding results for *Alexandrium* detection were obtained using the 18S-dinoflagellate amplicon, closely followed by 28S. Both amplicons provided comparable results with a very high percentage of total *Alexandrium* reads per sample for all five of the culture samples (>90% of reads on average) and a low percentage in the mixed culture (<4%). All six of these samples had highly positive qPCR results using an assay targeting the toxic North American strain of *A. tamarense* (Toebe et al. 2013).

Detections of *Alexandrium* in samples where it was not identified based on microscopy were less conclusive. There were 19 samples where *Alexandrium* presence was suggested by 18S-dinoflagellate and 28S amplicons but not by microscopy. In all cases these were very low counts of *Alexandrium* (*i.e.* very low percent of reads per sample, ranging from 0.02% to 4.72%), 13 of these 19 samples also had positive qPCR results, suggesting that the metabarcoding positives were accurate. Future work is needed to clarify exactly which results are correct, the microscopy or the molecular (qPCR and metabarcoding) for this taxon, and how to best score these detections (e.g. weak positives). With the expected higher sensitivity of molecular assays as compared to microscopy, it is not implausible that low copy numbers of a HAB species may exist in many mixed cultures, but whether these were introduced by cross-contamination of cultures, as well as the biological relevance of such levels also would need further study.

While *Alexandrium* presence detected by 18S-dinoflagellate and 28S amplicons was similar, the biggest difference in these amplicons was in the taxonomic level of identification. In the 18S-dinoflagellate amplicon, more than 99.50% of *Alexandrium* OTUs were identified to genus level only, while the very small remainder was assigned to *A. tamarense* (0.46%) and *A. andersonii* (<0.01%). In contrast, the 28S results indicated that 97.31% of total *Alexandrium* reads belonged to the species *A. fundyense*, 0.05% to *A. tamarense*, with the remainder 2.64% assigned to the genus level. While for decades *A. catenella*, *A. tamarense*, *A. fundyense* have been referred to as belonging to the “*Alexandrium tamarense* species complex” (Balech 1985), phylogenetic studies discriminated five distinctive ribotypes (Lilly et al. 2007; John et al. 2014) within the complex that do not correlate with the original morphospecies description. After much debate, *A. fundyense* was declared invalid (Prud’Homme Van Reine 2017) and nomenclature priority given to *A. catenella.* In our study, clonal cultures that were morphologically identified as *A. catenella*, *A. tamarense*, and *Alexandrium* spp. were all assigned to *A. tamarense* by TaqMan (Toebe et al., 2013 primer targets Group 1 genotype) and to *A. tamarense* or *A. fundyense* species by metabarcoding, depending on the amplicon. These results indicate that (i) all these cultures belong to the Group 1 genotype and should be referred to as *A. catenella*; and (ii) sequence databases (e.g. GenBank) need to be updated for this taxon.

#### 4.3.2. Cochlodinium

Two out of approximately 40 known *Cochlodinium* species (*i.e. C. polykrikoides* and *C*. *fulvescens*) form HABs and can cause fish kills (Kudela and Gobler 2012). In BC, blooms of *C*. *fulvescens* were implicated in farmed salmon kills that caused ~1.5 M USD in economic losses (Whyte et al. 2001). Here, *Cochlodinium* was observed by microscopy in two field samples. Its presence (24 and 992 reads) was detected at the genus level in both of these field samples, but only by the 28S amplicon. This identification was conclusive and OTUs of *Cochlodinium* at the species or genus level were not observed in any of the other 44 samples.

#### 4.3.3. Protoceratium reticulatum

*Protoceratium reticulatum* produces yessotoxins, which can be bioaccumulated by shellfish and have been associated with diarrhetic shellfish poisoning (DSP) (Satake et al. 1998). Metabarcoding was very effective for identification of this species. The sample of *P. reticulatum* from an established culture with identification based on morphology had a positive identification to the species level when using both the 18S-dinoflagellate and 28S amplicons (143,019, 42,069 reads). The 18S-dinoflagellate amplicon also detected very low presence (<2% of the total reads per sample) of *P. reticulatum* in another three culture samples and one field sample (s34). The 28S amplicon had comparable readings and percentages to the 18S-dinoflagellate amplicon.

#### 4.3.4. Dinophysis, Karlodinium, and Phalacroma

Algae from the *Dinophysis*, *Karlodinium*, or *Phalacroma* genera were not identified in the samples using microscopy, but were detected using molecular methods. Certain *Dinophysis* and *Phalacroma* species produce toxins that cause DSP, and thus are important to shellfish aquaculture. In BC, the first reported DSP outbreak was associated with elevated numbers of *Dinophysis* spp. (Esenkulova and Haigh 2012). In the present study, a low number of reads (<200 in total) of *D. parvula*, *P. rapa* (previously known as *D. rapa*), as well as *Dinophysis* spp. were detected in one field sample using the 28S amplicon. The *Karlodinium* genus includes several toxin-producing species; for example, *K. veneficum* blooms have been associated with aquatic faunal mortalities for decades (Place et al. 2012). A small number of reads to the genus and species level (<100 in total) were detected in one field sample by 28S. Similar results for both *Dinophysis* and *Karlodinium* were obtained using the 18S-dinoflagellate amplicon, but were only annotated to the genus level.

### 4.4. Raphidophytes

Raphidophytes are a group of algae with very few species. However, these species include several taxa that pose some of the most serious threats to finfish aquaculture around the world, such as *H. akashiwo* and those in the *Chattonella* genus (Hallegraeff 2004). In our work, the best results for raphidophytes were obtained using 18S-diatom and 28S amplicons.

#### 4.4.1. Heterosigma akashiwo

*Heterosigma akashiwo* is a major fish killer in BC, causing economic losses to the BC salmon aquaculture industry of about ~3.5 M USD per year (Haigh and Esenkulova 2014). All 24 samples (18 cultures and 6 field) where *H. akashiwo* presence was observed with microscopy returned positive detections with qPCR and with three of the amplicons. The amplicon with the weakest detection ability was the 18S-dinoflagellate, which identified *H. akashiwo* in only 19 out of 24 samples. The highest metabarcoding reads were detected by the 18S-diatom amplicon. In the rest of the samples (n=24), *H. akashiwo* presence was not noted by microscopy observations, but *H. akashiwo* reads were found in more than 20 of these samples by the 18S-diatom, 16S, and 28S amplicons, in six using the 18S-dinoflagellate amplicon, and in 11 of these samples by qPCR. For most of the samples there was a general agreement between *H. akashiwo* read numbers by the different techniques (e.g. all 18 samples where *H. akashiwo* was not detected by 18S-dinoflagellate had <100 reads per sample by all other techniques; most of the 15 samples with the highest *H. akashiwo* reads detected by 18S-diatom had the highest levels with other techniques). One exception to this trend was in the purchased culture of *Chattonella subsalsa*, where a high load of *H*. *akashiwo* was identified solely by the 16S amplicon. *C. subsalsa* identification in this case was confirmed by microscopy and other amplicons, so this clearly indicates a 16S database issue in GenBank (*i.e.* mis-representation of *C. subsalsa* species as *H. akashiwo*). In 35 samples where *H. akashiwo* presence was suggested based on qPCR results, there was a general agreement of higher loads (e.g. Ct<20) with high OTU read counts (>4,000) in the 18S-diatom amplicon, but not in other amplicons. However, some samples with lower qPCR loads (CT=20–29.5) also had very high OTU read counts, with nine samples containing over 10,000 reads. Therefore the quantitation between the read counts and the qPCR was not always congruent. It is possible that this could be due to mismatches in specificity for the applied assay, but further work would be needed to better understand this difference.

#### 4.4.2. Chattonella

Globally, many species of *Chattonella* have been associated with fish kills (Moestrup et al. 2008) and blooms of *Chattonella* sp. have caused farmed fish mortalities in BC (Haigh and Esenkulova 2014). Here, all ten local samples with positive microscopy identification of *Chattonella* spp. did not return results for this genus by sequencing. Almost all (>99.99%) of the Raphidophytes reads in these ten samples (*i.e.* one culture, five mixed cultures, and four field samples) where *Chattonella* presence was suspected based on microscopy, were assigned to *H. akashiwo* by all four amplicons. It is possible that large *Heterosigma* cells were misidentified as *Chattonella* during microscopy. The purchased culture of *Chattonella subsalsa* was positively confirmed as *Chattonella* (>99% of total reads) by 18S-diatom and 28S amplicons with a very small portion (0.04%) identified to the species level (*C. subsalsa*) using the 28S amplicon, suggesting that the amplicons could identify *Chattonella* if it was present. Therefore, for this species the NGS approach appeared to be superior to light microscopy identification.

A very low number of reads (<30) of *Chattonella* spp. were also detected in four culture samples of *Alexandrium* spp. and *H. akashiwo* by the 18S-diatom amplicon. This small number is most likely also an artefact of the subsampling process (*see above*). Only a small portion of the purchased *C. subsalsa* culture was identified to the species level, and the rest of the non-purchased samples was identified to genus only, which suggests the difference in resolution may be a result of species sequence variation that was not reflected in the NCBI database. TaqMan assays for *C. subsalsa* and *C. marina/ovata/antiqua* did not provide positive results with any of the samples.

### 4.5. Silicoflagellates (Dictyochales)

Silicoflagellates are a small group of algae, many of which possess a siliceous skeleton at certain stages of their life cycle. Some species from this group cause fish kills (Henriksen et al. 1993), including farmed salmon mortality events in BC (Haigh and Esenkulova, 2014; Haigh et al., 2014, Haigh et al. 2019). All four amplicons underperformed in detecting algal species and genera from the silicoflagellate group. Most of the samples (11 out of 15) where taxa from this group were observed by microscopy did not have reads annotated as silicoflagellates. The definitive reason for poor metabarcoding identification performance to this target remains unknown, however it could be related to the extraction process, to the primers or amplification process, or due to incomplete databases.

#### 4.5.1. Dictyocha

Skeleton-containing cells of *D. speculum* are very easily identified by microscopy based on their unique shape and size. Blooms of this species have been linked with fish kills around the world (e.g. Henriksen et al. 1993) including in BC (Haigh and Esenkulova 2014; Haigh et al. 2019). In our study, six field samples contained *D. speculum* cells, based on microscopy observations, but there were no reads assigned to the *Dictyocha* species, genus, or family by any of the NGS amplicons. One of these samples had a very low number of reads (*i.e.* 12 reads) detected by the 18S-diatom amplicon, but was assigned only to the order level. These results show that with the current sequencing database, the NGS approach can miss targets at the species, genus, and even family levels. Further work is needed to improve the sequencing database with sequences from voucher specimens to determine which assays are capable of differentiating this genus, or species within this genus, from other closely related taxa.

#### 4.5.2. Pseudochattonella verruculosa

Both species in the *Pseudochattonella* genus (*i.e. P. farcimen* and *P. verruculosa*) have been implicated in wild and farmed fish kills (e.g. Jakobsen et al. 2012). In BC, *Pseudochattonella* sp. blooms have been associated with farmed fish mortalities since 2007 (HAMP, unpublished data). For this project, one culture from a fish-killing event (Haigh et al. 2014) was established and five field samples were collected. High and comparable loads (>2,000 reads) of *Pseudochattonella* genus were detected in the culture sample by 18S-diatom and 18S-dinoflagellate amplicons. For the former, ~1% of the total reads were assigned to the species level for *P. farcimen*. This culture sample was also the only one that provided qPCR amplification results (Ct=14) with the *P. verruculosa* assay. A very low number of reads (<20 reads) of *Pseudochattonella* were detected in one of the field samples, where its presence was indicated based on microscopy and by 18S-diatom and 18S-dinoflagellate runs, but not by qPCR. This is the first published confirmation of both qPCR and metabarcoding identification of *Pseudochattonella* linked to a fish-killing event in Canada.

#### 4.5.3. Pseudopedinella

Toxicity of *P. pyriformis* (previously known as *P. pyriforme*) was recently discovered in laboratory studies (Skjelbred et al. 2011). Based on microscopic identification, this species has been linked to farmed salmon mortality in BC (HAMP, unpublished data). In this work, one culture and two field samples were available for molecular analysis. All these samples were obtained from a fish-killing event on the west coast of Vancouver Island. To our knowledge, this is the first published report of ichthyotoxic effects of *Pseudopedinella* species in the field. Metabarcoding did not detect *Pseudopedinella* in the available samples, and it did not identify any reads from the silicoflagellates taxon. The majority of reads in the culture of suspected *Pseudopedinella* sp. was the diatom *Plagiostriata goreensis* by the 18S-diatom amplicon (98%) and “not assigned” in the 18S-dinoflagellate and 28S amplicons (94%, 99%, respectively). While there was therefore no molecular confirmation of the microscopic identification of *Pseudopedinella* in the fish-kill-related sample, the established culture was certainly not a diatom and contained *Pseudopedinella*-like cells (small flagellates). To improve this detection method, a positive control would need to be included, such as a purchased *Pseudopedinella* culture, in order to ensure that the selected amplicons can identify this genus. However, given the putative misidentification of the culture of suspected *Pseudopedinella*, it is also possible that this is a database issue.

### 4.6. Haptophytes

Toxic haptophytes in the *Prymnesium* and *Chrysochromulina* genera are known to cause fish mortality (Moestrup 2003). In our work, the best results were obtained using the 18S-dinoflagellate and 28S amplicons.

#### 4.6.1. Chrysochromulina

Haptophytes may be identified with light microscopy to the genus level, but this is challenging due to small cell size, and species level identification is only possible with scanning electron microscopy. In BC, blooms of suspected *Chrysochromulina* spp. have been linked to fish kills on salmon farms since the year 2000 (HAMP, unpublished data). Our study had four field samples where *Chrysochromulina* presence was suspected based on microscopy, but two of these samples had low DNA yields and resulted in <200 total reads per sample. The 18S-dinoflagellate amplicon detected *Haptolina fragaria* (>100 reads) in the other two samples. *Haptolina fragaria* (presumably a non-toxic Prymnesiales species), originally described as *Chrysochromulina fragaria* (Eikrem and Edvardsen 1999) but later described to a new genus from ribosomal DNA phylogenetics (Edvardsen et al. 2011). Low levels of *Chrysochromulina* spp. were also detected by the 28S amplicon (220 reads, 0.35%) in one field sample where its presence was not indicated based on microscopy. With the importance of haptophyte species in fish-kills, and the difficulty in microscopic identification of these species, this is an area where better detection by molecular methods would be of strong practical use.

### 4.7. Cyanobacteria

Cyanobacteria, or blue-green algae, are highly diverse aquatic bacteria with over 2,000 species (Nabout et al. 2013). More than 55 of them have been shown to produce toxins that are harmful to humans, as well as other terrestrial and aquatic life (Cronberg, 2003). Due to their generally small size and limited significance to marine finfish, cyanobacteria are not recorded and identified during routine harmful algae analysis in BC by HAMP, unless they appear to dominate in the sample. Here, only one sample (mixed culture) had evidence of possible cyanobacteria presence (based on microscopy), and was mixed with green algae. The cyanobacteria presence in this sample was not confirmed by molecular techniques, but cyanobacteria were detected in other samples by molecular techniques where its presence was not described based on microscopy. Nine samples provided amplification results with qPCR (Ct=17–29), but only one of these samples had evidence for cyanobacteria presence based on metabarcoding, as found using the 16S amplicon (Synechococcaceae, 101 reads, 0.1%). Whether the qPCR results are species specific, or whether the 16S metabarcoding was missing these detections remains unknown, and requires further study.

### 4.8. Summary of Methods Evaluation

Development of molecular methods for harmful taxa detection and abundance estimation is an active area of research. The present study allowed for an evaluation and cross validation of traditional microscopy and current molecular techniques and assays for each taxon relevant to BC HABs (Table 5). The most appropriate molecular technique, as determined by highest detection rate using currently available assays was identified based on the results such as taxon resolution, the conclusiveness of the detection, and whether it was confirmatory of microscopy results. Molecular techniques, and metabarcoding in particular, proved to be a very promising complement to standard microscopy, and in some cases potentially an alternative given additional benchmarking. However, there is still much work to be done to develop a curated database with voucher specimens to ensure adequate representation and detectability of all of the important HABs forming species. The comprehensive nature of the metabarcoding approach is another benefit, where many species can be simultaneously detected even from different taxonomic levels. Information obtained during this research provides a foundation to build upon, for example, by using the sequencing results to identify species presence, to develop specific qPCR primers, and to identify existing gaps in sequence databases.

**Table 5.**
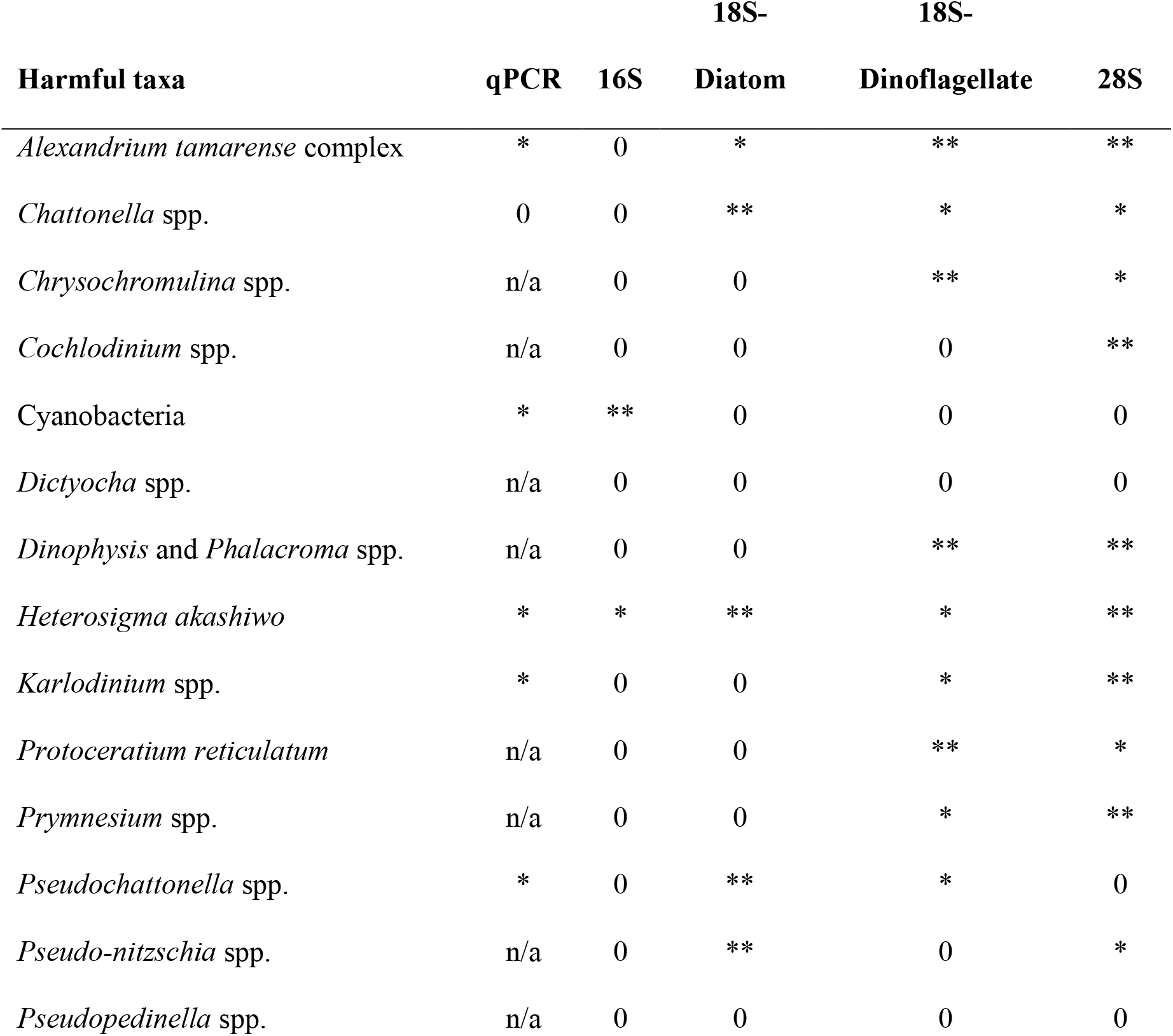
Evaluation of harmful taxa detection methods based on the present study: ** = appropriate current technique(s) for taxa detection (at least one designated per taxon), * = technique that can detect taxa, 0 = no taxa detection, n/a = not applied.

Quantitative PCR is a cost effective approach and can be used for presence/absence as well as quantitative estimates of targeted species, however it requires existing knowledge of potential HABs taxa that may be present in monitored samples. In our study, existing (as of 2014) TaqMan assays were found for only four out of 14 targeted taxa. Ten taxa that were identified by microscopy did not have a molecular assay available during the time of qPCR run of this study, including *Chrysochromulina* spp., *Cochlodinium fulvescens*, *C. concavicornis*, *C*. *convolutus, Dictyocha* spp., *Dinophysis* spp., *K. mikimotoi*, *Protoceratium reticulatum*, and *Pseudo-nitzschia* spp. These particular taxa are an important priority for assay development for application in BC. Methodological weaknesses in the present study, such as the absence of positive controls and PCR occurring under thermal regimes different than those under which the assay was designed, limit the interpretation of qPCR results. Despite of these limitations, assays for detection of *A. tamarense* (A.tam1), *H. akashiwo* (H.aka1), and *P. verruculosa* (P.ver1) performed well (i.e., the results were comparable to microscopy and metabarcoding) and could be implemented in routine monitoring. Sequencing data acquired during this work provides the ability to develop tailored qPCR assays in future work.

Metabarcoding has been used in the detection and assessment of phytoplankton communities and harmful algae around the world (e.g. Abad et al. 2016; Akita et al. 2019; Le Bescot et al. 2016; Liu et al. 2020; Smith et al. 2017) establishing next generation sequencing as a powerful emerging approach in plankton monitoring. In the present study, metabarcoding enabled accurate identification of multiple species and was superior in some cases to light microscopy, for example it was able to identify cryptic species not possible to differentiate to the species level by light microscopy (e.g. *Chrysochromulina*, *Prymnesium*). It generated high-throughput data, providing community composition diversity information for the samples, and capturing some potentially harmful taxa that were not originally targeted, as well as providing some cursory information on relative abundance, although the reliability of abundance estimates has not yet been conducted. The presence of a large number of unannotated reads in a number of samples suggests that the sequences of a number of species are still missing from the public databases. This will improve with curated databases, which will be generated as this method and application continues to be developed.

The highly sensitive nature of this technique emphasised the need for extremely careful sample handling, as detections of pure cultures transferring small proportions of taxa between samples was detected, probably caused during subsampling of cultures. Since the laboratory component of this study was conducted, the increased use of metabarcoding in many areas of research (e.g. environmental DNA, or eDNA), has resulted in specialized sample collection, processing workflows, laboratory environments, and establishing threshold detections designed to reduce contamination risks. Although metabarcoding processing time and costs may limit uptake currently, with increased development it is highly likely to become a useful technique for analysing samples collected during active HABs, leading to an unprecedented monitoring opportunity of HABs in terms of sensitivity and precision.

## 5. Conclusion

Applying light microscopy and various molecular techniques to culture and field samples containing multiple harmful algal species allowed cross validation of these techniques and offered a significant foundation for choosing appropriate techniques for targeted taxa in the future. DNA yields were considerably higher for the cultures and filtered field samples, whereas pelleted samples were often unusable. While TaqMan assays were available only for four out of 14 HAB taxa of concern in BC, assays for detection of *A. tamarense*, *H. akashiwo*, and *P. verruculosa* provided adequate identification results. This indicates a need for the development of primers and probes for the rest of the harmful species to allow cost effective detection of many species simultaneously. Sequencing data obtained during this study will enable the development of new qPCR assays tailored for species within the northeastern Pacific Ocean.

Metabarcoding with a combination of markers (*i.e.* 16S, 18S-diatom, 18S-dinoflagellate, and 28S) allowed the identification of over 350 taxa and proved to be an unmatched technique for phytoplankton community structure analysis. Different markers had different strengths for particular taxa, although result congruence was observed among amplicons. The 18S-diatom amplicon identified harmful taxa from the diatom and raphidophyte groups, and the 18S-dinoflagellate and 28S amplicons, producing similar results to each other, overall provided the best identification for harmful algae from dinoflagellate and haptophytes groups, as well as the raphidophyte *H. akashiwo*. Although cyanobacteria were detected by only the 16S amplicon, this amplicon did not perform well for most of the HABs species. All of the amplicons underperformed for identification of silicoflagellate algae, although the reason for this remains unknown. The isolated culture from the fish-killing event associated with a *P. verruculosa* bloom (morphology-based identification) was confirmed as *P. verruculosa* by qPCR and *Pseudochattonella* spp. by NGS, thus producing the first record of PCR and metabarcoding confirmation of *Pseudochattonella* associated with a fish kill in Canada. Overall, the combination of morphology and molecular-based identification, if implemented, will greatly improve HABs monitoring, help mitigate issues caused by HABs, and aid in better understanding the dynamics of the phenomenon. This work demonstrates a pressing need to tailor qPCR assays, to improve reference databases, and to apply a multiple marker approach for metabarcoding of diverse taxa.

## Acknowledgments

This project was supported by the Aquaculture Collaborative Research and Development Program (ACRDP; Project number: P-12-01-003) of Fisheries and Oceans Canada and the British Columbia aquaculture industry including Creative Salmon Company Ltd., Grieg Seafood BC Ltd., Mainstream Canada (now Cermaq Canada), Marine Harvest Canada Inc. (now Mowi Canada West), Cleanwater Shellfish Ltd., Island Scallops Ltd., Little Wing Oysters Ltd., Mac’s Oysters Ltd., Nelson Island Sea Farms Ltd., and Taylor Shellfish Canada ULC. Yves Perreault of Little Wing Oysters Ltd. and Dave Guhl of Marine Harvest Canada Inc. are acknowledged for helping with sediment sampling at Okeover Inlet and Quatsino Sound, respectively. Crew of the Deep Bay Marine Station enabled sediment sampling in Baynes Sound. Laurie Keddy (Pacific Biological Station, DFO) is thanked for maintaining established cultures and providing advice on isolation and culturing.

## Data Availability

The pipeline for data analysis is available on GitHub: https://github.com/bensutherland/eDNA_metabarcoding

Raw sequence was uploaded to SRA under BioProject PRJNA544881 within BioSample accessions SAMN11865982-SAMN11866125.

## Supplemental Information

**Table S1.** Overview of the algal taxa and TaqMan qPCR primers as well as metabarcoding primers used in this study.

**Table S2.** TaqMan assay amplification results: “+” positive (Ct<20) and “?” suspected (Ct 20-30). Note: Full name for the primer abbreviation and its citation are listed in Table S1. Note that Ct’s on the microfluidics BioMark platform are approximately 10 Ct values lower than on traditional single-assay platforms (Miller et al. 2016).

**Table S3.** Read counts per taxonomic group using the 16S amplicon.

**Table S4.** Read counts per taxonomic group using the 18S-diatom amplicon.

**Table S5.** Read counts per taxonomic group using the 18S-dinoflagellate amplicon.

**Table S6.** Read counts per taxonomic group using the 28S amplicon.

**Table S7.** Taxa detected by microscopy, the closest match as detected by NGS, the percentage of the total reads for that sample represented by the match, and the percentage of unknown reads in the sample. PC – purchased culture, IC – isolated culture, MC – mixed culture, FWS – filtered water sample, PWS – pelleted water sample; * - grouped; n/a - not applied

